# Elucidating the genetic basis of an oligogenic birth defect using whole genome sequence data in a non-model organism, *Bubalus bubalis*

**DOI:** 10.1101/060996

**Authors:** Lynsey K. Whitacre, Jesse L. Hoff, Robert D. Schnabel, Sara Albarella, Francesca Ciotola, Vincenzo Peretti, Francesco Strozzi, Chiara Ferrandi, Luigi Rammuno, Tad S. Sonstegard, John L. Williams, Jeremy F. Taylor, Jared E. Decker

**Affiliations:** Informatics Institute, University of Missouri, Columbia, Missouri, USA; Division of Animal Sciences, University of Missouri, Columbia, Missouri, USA; Department of Veterinary Medicine and Animal Production, University of Naples Federico II, Naples, Italy; Parco Tecnologico Padano, Lodi, Italy; Department of Agriculture, University of Naples Federico II, Portici, Napoli, Italy; Recombinetics, St. Paul, Minnesota, USA; Davies Research Centre, School of Animal and Veterinary Sciences, University of Adelaide, Roseworthy, Australia

## Abstract

Recent strong selection for dairy traits in water buffalo has been associated with higher levels of inbreeding, leading to an increase in the prevalence of genetic diseases such as transverse hemimelia (TH), a congenital developmental abnormality characterized by the absence of a variable distal portion of the hindlimbs. The limited genomic resources available for water buffalo, in conjunction with an unconfirmed inheritance pattern, required an original approach to identify genetic variants associated with this disease. The genomes of 4 bilaterally affected cases, 7 unilaterally affected cases, and 14 controls were sequenced. Variant calling identified 19.8 million high confidence single nucleotide polymorphisms (SNPs) and 2.8 million insertions/deletions (INDELs). A concordance analysis of SNPs and INDELs requiring all unilateral and bilateral cases and none of the controls to be homozygous for the same allele, revealed two genes, *WNT7A* and *SMARCA4*, known to play a role in embryonic hindlimb development. Additionally, SNP alleles in *NOTCH1* and *RARB* were homozygous exclusively in the bilaterally affected cases, suggesting an oligogenic mode of inheritance. Homozygosity mapping by whole genome *de novo* assembly was then used to identify large contigs representing regions of homozygosity in the cases. This also supported an oligogenic mode of inheritance; implicating 13 genes involved in aberrant hindlimb development in the bilateral cases and 11 in the unilateral cases. A genome-wide association study (GWAS) predicted additional modifier genes. Results from these analyses suggest that mutations in *SMARCA4* and *WNT7A* are required for expression of TH, while several other loci including *NOTCH1* act as modifiers and increase the severity of the disease phenotype. Although our data show that the inheritance of TH is complex, we predict that homozygous variants in *WNT7A* and *SMARCA4* are necessary for the expression of TH and selection against these variants and avoidance of carrier-to-carrier matings should eradicate TH.

**Author Summary:** Genetic diseases often occur and are spread through small populations under strong selection where rates of inbreeding can be significant. The use of a limited number of water buffalo males via artificial insemination for genetic improvement of milk and milk composition has increased the frequency of the genetic disease, transverse hemimelia (TH). Transverse hemimelia affected calves are normally developed except for malformation of one or both hindlimbs or both hindlimbs and one or both forelimbs. Little is known about the inheritance pattern of TH. We discovered genetic variants present in cases where both hindlimbs and one forelimb were affected, cases were both hindlimbs were affected, cases where only one hindlimb was affected, and in non-affected water buffalo that predict TH to be inherited as an oligogenic disease with two driver loci necessary for disease expression and several additional modifier genes that are responsible for the severity of the disease phenotype. We predict that selection against mutations in the two major loci and the avoidance of mating animals that are heterozygous for these mutations will eliminate TH from water buffalo.

## Introduction

Water buffalo were domesticated approximately 5,000 years ago in Indian subcontinent [1]. Today, there are over 130 million domesticated water buffalo worldwide that serve as an important component of agriculture through both milk and meat production [2]. In many developing countries, water buffalo account for more than 50% of the milk production and are relied upon more than any other domesticated species [3,4]. Recently, a genetic disease called transverse hemimelia (TH) has appeared in Italian Mediterranean River buffalo, most likely as an indirect result of strong selection for dairy production traits and an accompanying increase in the rate of inbreeding. Transverse hemimelia causes unilateral or bilateral hindlimb malformation and is defined by the lack of development of distal hindlimb structures, which manifests as the loss of one or both hindlimbs at a distal point that is variable among cases (Figure 1A, 1B). In severe cases with bilateral hindlimb malformation, one or both forelimbs may also be affected. Involved limbs appear as to be amputated with the exception of the fact that there are sketches of claws in the terminal part [5]. The prevalence of the disease has been estimated to be between 2 and 5 percent in some populations of Mediterranean Italian River buffalo [6]. Unfortunately, because record keeping is poor and pedigrees are often unknown or incomplete, the mode of inheritance of TH in water buffalo has not been established.

**Figure 1.**
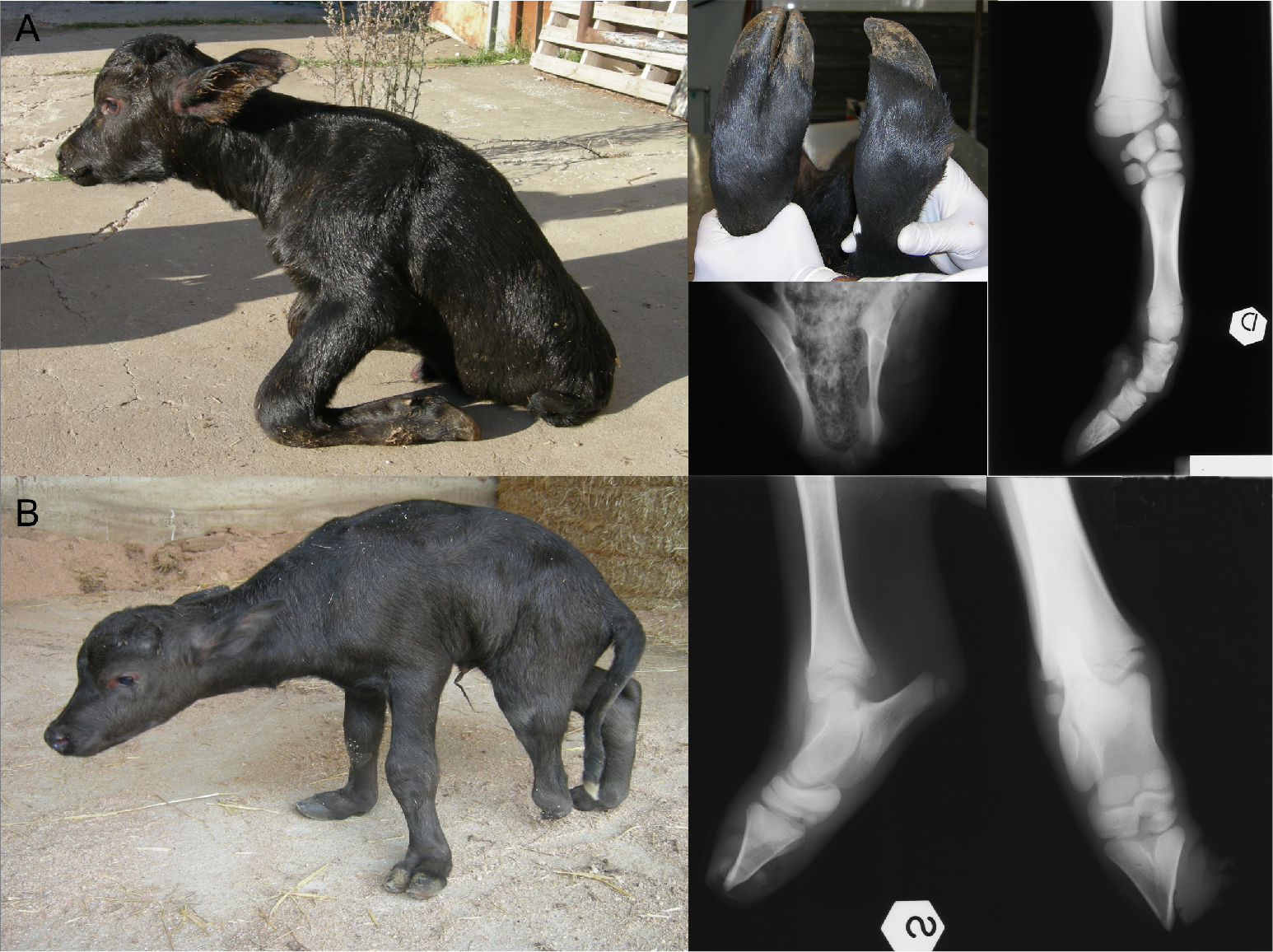
Water buffalo calves with TH. (A) A bilaterally affected TH case with both hindlimbs completely absent at birth and hypoplasia of carpal bones and absence of medial bones starting from metacarpus and X-ray images. (B) A unilaterally affected TH case with one hindlimb truncated at the tarsus with x-ray image.

Other types of hemimelia, a generalized developmental anomaly resulting in the absence of the distal portions of one or more limbs, have been reported in domestic species including goats, lambs, cattle, dogs, and cats [7–11]. In goats, lambs, and cattle, hemimelia has been shown to be heritable, but it can also be caused by environmental exposures to teratogenic plants, parasites, and drugs [12]. The increase in hemimelia in livestock species has been blamed on high levels of inbreeding due to selection for economically valuable traits resulting in increased homozygosity of recessive deleterious mutations. Pedigree analyses of dogs and Shorthorn cattle have revealed hemimelia to be inherited as an autosomal recessive disorder [9,10]. Hemimelia has also been reported in humans, but occurs either due to autosomal recessive inheritance or sporadically, suggesting a polygenic mode of inheritance [13].

Despite several recent research efforts to elucidate the molecular mechanisms involved in hemimelia, the causal mutations in water buffalo and many other species are currently unknown. However, the genetic mechanisms responsible for embryonic hindlimb morphogenesis have been extensively studied in model species and several genes have been implicated in hindlimb development [14–17]. Furthermore, many genes involved in embryonic morphogenesis have been suggested to play roles in the manifestation and inheritance of TH [18,19]. To elucidate the genes involved in the inheritance of TH in water buffalos, we sequenced 11 affected buffaloes (4 with bilateral TH and 7 with unilateral TH) and obtained sequences for 14 control buffaloes from the International Water Buffalo Genome Consortium. Our analyses of these data suggest an oligogenic inheritance pattern, and implicate variations in *SMARCA4* and *WNT7A* as the main drivers necessary for the manifestation of TH. The accumulation of homozygous mutations in modifier genes appears to impact the severity of the TH phenotype, resulting in animals that vary from the lack of a single transverse bone in one limb to the complete lack of both hindlimbs with malformation of one forelimb. The analyses leading to these conclusions describe a novel method for detecting the loci underlying an oligogenic diseases in a non-model organism that lack refined genomic resources such as a completed reference genome or annotated gene models.

## Results

### Alignment and variant calling

Alignment of DNA sequences from 11 cases and 14 controls to the UMD_CASPUR_WB_2.0 water buffalo reference assembly resulted in an average mapping rate of 99.17% and average coverage of 9.15X (Table S1). Initial variant calling identified approximately 21.7 million SNPs and 2.8 million INDELs. After filtering on quality 19.8 million SNPs and 2.7 million INDELs remained for analysis. The overall genotype call rate was 98.02% in the cases and 90.99% in the controls; consistent with the reduced depth of sequence coverage and older technology used to sequence the controls (Table S2).

### Case versus control concordance analysis

SNP and INDEL concordance analyses were performed to identify variants for which all cases were homozygous for an allele that was never homozygous in the controls. In total, 1,741 SNPs and 793 INDELs met these criteria (Table S3, S4). Nine hundred seventy-one of the SNPs were not in genes, but the 770 remaining SNPs were found in 451 unique genes. Two of the genes, *SMARCA4* and *WNT7A*, were associated with the GO term “embryonic hindlimb morphogenesis.” Furthermore, when only the bilaterally affected cases were analyzed for SNP concordance two additional genes, *NOTCH1* and *RARB*, which are also associated with embryonic hindlimb morphogenesis, were detected. When only the three most severely affected bilateral cases, with the complete loss of both hindlimbs, were analyzed one additional hindlimb morphogenesis related gene, *TFAP2B*, was detected. These findings, along with the fact that no additional associated genes were detected when the unilaterally affected cases were analyzed, present the first genomic evidence for an oligogenic mode of inheritance for TH in water buffalo. However, by aligning the available cattle gene models for these genes to the water buffalo genome assembly, we predicted that all of the disease-associated mutations were located in introns.

Despite the challenge of calling INDEL genotypes with high accuracy from low coverage sequence data, we also analyzed the detected INDELs for their concordance with TH phenotype. Of the 793 concordant INDELs, 222 were located in genes, but only one was found in a gene associated with hindlimb morphogenesis. This INDEL was found in *WNT7A*, a gene also identified by the SNP analysis, and occurs as a 3 bp insertion (C -> CCCG). Based on aligning to the *WNT7A* gene annotation in bovine, this variant appears to be located in intron 3. Unlike the concordance analysis performed for SNPs, no additional concordant INDELs were detected as the cases were further restricted according to the severity of the TH phenotype. We interpret the results of the INDEL analysis cautiously because, even after filtering for quality, a large proportion of the remaining INDELs appear to have been identified because of homopolymer repeat errors.

### Homozygosity mapping by de novo assembly

Large runs of homozygosity (ROH) are common in inbred animals, which have an increased risk for genetic diseases because the deleterious effects of recessive alleles are expressed when they are found in a homozygous state. However, with the exception of selective sweep regions that affect all animals, runs of homozygosity should not be shared across large numbers of unrelated individuals. Thus, homozygosity mapping is a powerful method to identify loci responsible for autosomal recessive traits [20]. Assuming a common origin for all cases, ROH should capture the loci that cause TH in distantly related affected animals (see Methods). However, because the water buffalo reference assembly currently exists as an early draft with more than 367,000 unplaced sequence scaffolds, we used a novel approach for homozygosity mapping that was not limited by the reference assembly scaffold lengths.

Three whole genome *de novo* assemblies were performed from the sequence data, which were pooled separately from four bilaterally affected cases, four unilaterally affected cases, and four controls. Regions of the genome with lower heterozygosity in sequence reads pooled from multiple individuals should be assembled into longer contigs, due to reducing forking in the assembly graph. Overall, the contig N50 statistics achieved for the bilaterally affected and unilaterally affected cases were much larger than for the controls (Table S5) and contained contigs that were approximately 0.75 orders of magnitude larger than those assembled for the controls (Figure 2). We were also able to assemble nearly the entire water buffalo genome (~2.64 Gb) in 218,053 contigs in the bilaterally affected cases and in 221,020 contigs in the unilaterally affected cases, both significantly fewer than for the current reference assembly, compared to 541,203 contigs for the controls. Estimated from a negative binomial generalized linear model, the mean contig lengths of the bilateral (12,140.04 bp) and unilateral (11,957.47 bp) assemblies were significantly longer than the mean contig length from the assembly of the control samples (4,738.88 bp), which is most likely due to the higher coverage for these samples (Table S5). Further, the mean contig length from the assembly of the bilateral cases was significantly longer than from the assembly of the unilateral cases (Z-score = -4.08, *p*-value = 4.6e-05). Overall, these results clearly indicate an increase in genome-wide homozygosity in the TH affected buffaloes.

**Figure 2.**
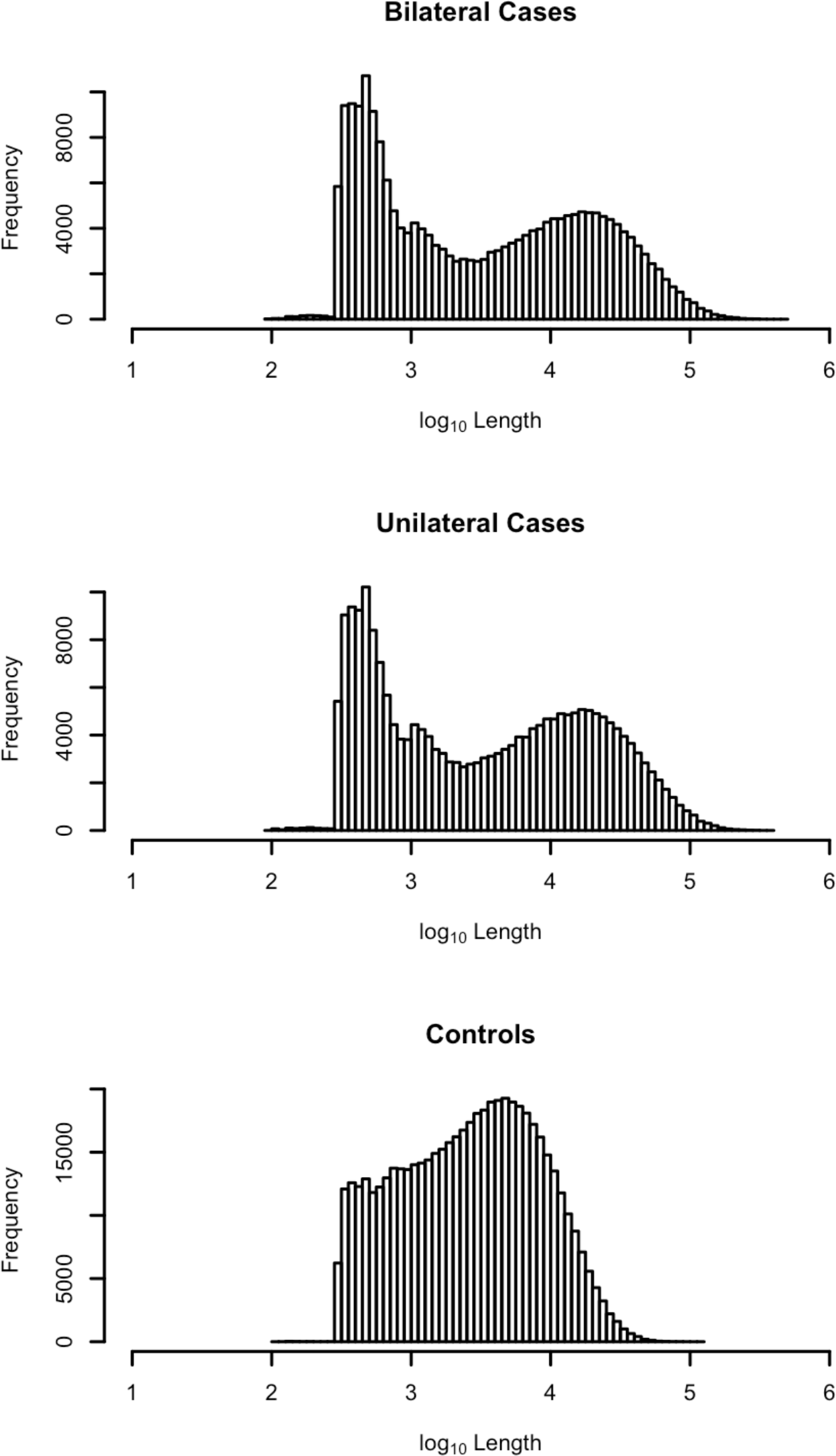
Log_10_-transformed distribution of contig sizes from the *de novo* assembly of pooled sequences from the bilaterally affected cases, unilaterally affected cases and controls.

We annotated the gene content of contigs from each assembly that were significantly larger than average by aligning them to the water buffalo reference genome. Following FDR correction, the assembly produced for the controls had 354 contigs significantly larger than the average, the assembly for the unilateral cases had 194 contigs significantly larger than average, and the assembly for the bilateral cases had 365 contigs significantly larger than average. The large contigs identified following FDR correction contained 5, 2, and 0 hindlimb morphogenesis genes for the bilateral cases, the unilateral cases and the controls, respectively (Table S6). None of these genes were in common with those detected from the SNP and INDEL concordance analyses.

However, homozygosity mapping by *de novo* assembly is confounded by repetitive elements that are difficult to assemble across. The distribution of these elements in the water buffalo genome is unknown as the reference assembly is not of high quality. Therefore, the effect of repetitive elements on disrupting the assembly of long contigs cannot be assessed. To compensate for this and include regions that may be largely homozygous but poorly assembled, contigs in the 99^th^ percentile for size from each of the assemblies were also aligned to the water buffalo reference genome to assess their gene content. Analysis of the longest 1% of contigs assembled for the controls, unilaterally and bilaterally affected cases revealed 2, 11, and 13 genes associated with hindlimb morphogenesis, respectively (Table S7). Six of these genes – *SMARCA4, NOTCH1, CHD7, MSX1, SALL1*, and *TBX3* – were detected in both the bilaterally affected and unilaterally affected cases. The *SMARCA4* locus, which was also identified from the SNP and INDEL analyses, was of particular interest because this gene and its flanking regions were assembled into a single contiguous sequence approximately 140 kb in length in both the bilaterally and unilaterally affected cases, but in the controls was placed on over 40 disjoint contigs (Figure S1).

### Genome-wide association study

Association analyses using both binary phenotypes, coding affected *versus* controls, and semi-quantitative phenotypes, which took into account disease severity, were used to discover additional candidate loci (Figure 3). While the binary trait GWAS primarily identified regions on small contigs with no nearby genes, *CHAMP1* and three uncharacterized predicted coding regions were detected (Table S8). *CHAMP1* was also detected by homozygosity mapping analysis and was on a contig significantly larger than average in both bilateral and unilateral cases, however, no concordant SNPs or INDELs were identified. GWAS using semi-quantitative phenotypes revealed 15 additional significantly associated genes. These included *FZD4*, a Wnt receptor, and *FGFR1*, a fibroblast growth factor receptor (Table S9). The GWAS results also suggest an oligogenic mode of inheritance because numerous loci rise to the same level of significance, in contrast to a GWAS for a Mendelian trait, where one primary peak is detected in a large sample case *versus* control analysis.

**Figure 3.**
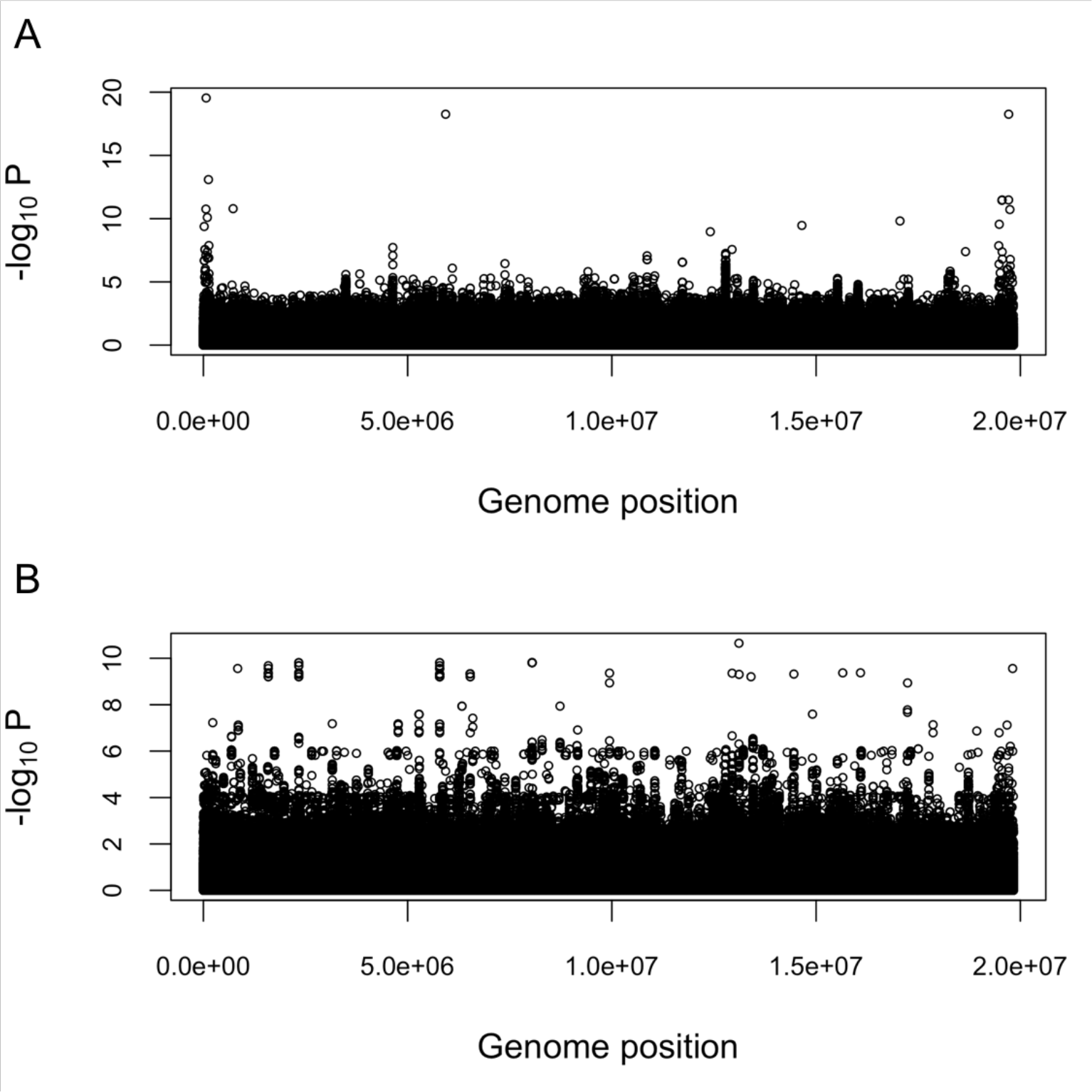
Manhattan plots of GWAS results. (A) GWAS results from association with a binary phenotype. (B) GWAS results from association with a semi-quantitative phenotype.

Although the GWAS failed to identify any genes related to hindlimb morphogenesis, several potential modifier genes were detected. This may be due to the nature of the GWAS, which assumed an additive model underlying the severity of the phenotype. The concordance and homozygosity mapping analyses, however, suggested that the phenotype is influenced by epistatic interactions among driver and modifier genes. A further limitation of the GWAS was that SNPs with one or more missing genotypes were either ignored by the analysis algorithm or had association effects estimated by assigning the mean allele frequency to missing genotypes. This was particularly problematic here, because of the higher rate of missing genotypes in the controls *versus* cases, due to the lower sequencing depth. Consequently, we filtered results for loci with one or more missing genotypes, resulting in only about 15% of loci being analyzed for association with the TH phenotypes (Table S10). Furthermore, although we were able to identify several loci at genome-wide significance (using a Bonferroni correction), we recognize that the sample size is not ideal for GWAS.

### Candidate region mapping

To overcome the challenge of relating significant associations with TH from the variety of performed analyses and the fact that the water buffalo scaffolds are not assigned to chromosomes, we mapped all of the buffalo genomic regions containing candidate loci to the UMD3.1 bovine reference assembly. While this identified candidate regions on all 29 bovine chromosomes, several chromosomal regions were identified as being significantly associated with TH in all performed analyses (Figure S2). This again suggests that loci involved in determining TH are spread throughout the buffalo genome and are not concentrated in a single region as would be expected if TH was inherited as a simple Mendelian phenotype.

Rudimentary gene enrichment analyses from the mapping of candidate regions to the bovine reference genome also indicated oligogenicity. From the SNP concordance mapping, a total of 769 candidate genes were discovered. This corresponds to approximately 3.85% of the total number of annotated genes in the genome (http://useast.ensembl.org/Bostaurus/Info/Annotation?redirect=no) and 6.06% of the total number of hindlimb morphogenesis genes. When only the bilaterally affected cases were analyzed for SNP concordance, two additional hindlimb morphogenesis genes were detected, but when only the unilaterally affected cases were analyzed no additional genes were associated with the GO term. Similarly, from the homozygosity mapping by *de novo* assembly analyses, contigs in the 99^th^ percentile for size from the unilaterally affected cases covered 20.35% of all annotated bovine genes and 33.33% of all hindlimb morphogenesis genes. Large, homozygous contigs assembled from the pooled sequences from the bilaterally affected cases covered 21.36% of all annotated bovine genes but covered 39.39% of the hindlimb morphogenesis genes. Contigs assembled from the sequences for controls covered 9.70% of all annotated bovine genes but only 6.06% of the hindlimb morphogenesis genes. These results consistently demonstrate an enrichment of hindlimb morphogenesis genes in the cases compared to the controls with even further enrichment in the bilaterally versus unilaterally affected cases and validate the oligogenic mode of inheritance of TH in water buffalo.

### Gene Ontology Enrichment

Genes identified by the various analyses were investigated for enriched gene ontology terms. While the term embryonic hindlimb morphogenesis was not significantly enriched in any of the analyses, the collective list of genes identified by SNP concordance, GWAS, and homozygosity mapping was enriched for embryo development (*p*-value = 0.0163), developmental processes (*p*-value = 7.04E-9), and anatomical structure development (*p*-value = 1.57E-9).

### Network analysis

Network analysis of all hindlimb morphogenesis genes detected from the concordance analysis and homozygosity mapping and all potential modifier genes identified from the GWAS was performed to understand how these genes interact. From these analyses, we identified 31 genes, of which 23 formed an exclusive network (Figure 4). In this network, *SMARCA4* interacts with 12 other genes while *WNT7A* interacts with 13.

**Figure 4.**
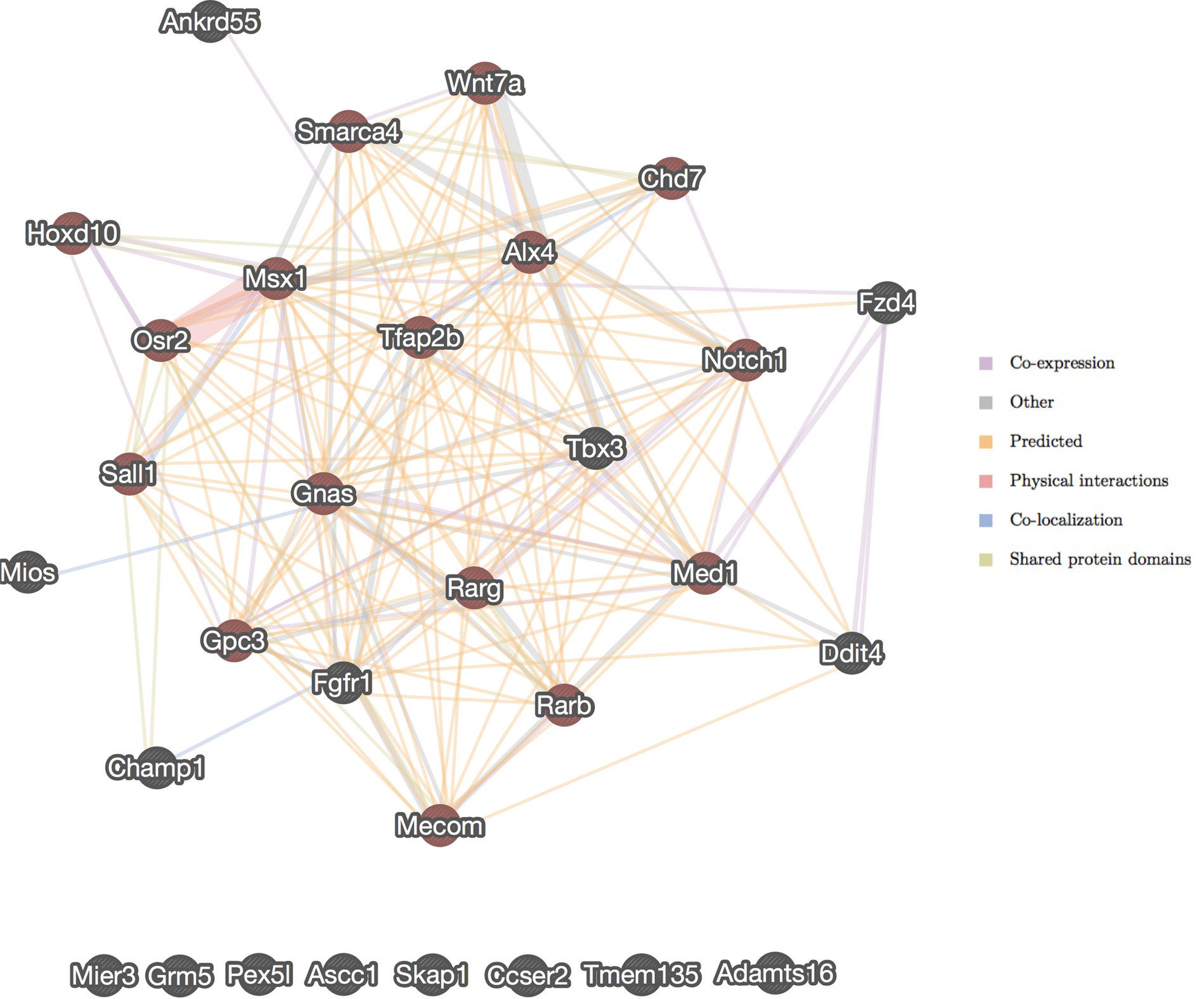
Network analysis of genes predicted to be associated with TH based on SNP concordance analysis, homozygosity mapping by *de novo* assembly, and GWAS. Genes associated with embryonic hindlimb morphogenesis are shaded in red.

## Discussion

We used several tactics to identify the genes involved in TH, a congenital limb abnormality resulting in the loss of transverse elements of the hindlimbs in water buffalo. Although the inheritance pattern of TH was initially unknown, we present evidence for an oligogenic inheritance and identify two main driver genes as well as several modifier genes. While mutations in both of the main driver genes, *SMARCA4* and *WNT7A*, appear to be necessary for the disease, mutations in the modifier genes contribute to the severity of the expressed phenotype. The *SMARCA4* chromatin remodeling factor is required for normal embryonic development and *SMARCA4* knockouts are lethal [21,22]. Furthermore, *SMARCA4* expression knockdowns in mice have a large effect on embryonic hindlimb and tail development [17]. These knockdowns possess a phenotype that is very similar to that of the TH affected water buffalo, where the development of the rest of the body and forelimbs is normal and the embryo is viable, but the hindlimbs are extremely underdeveloped.

Identification of two of the three known retinoic acid receptors (*RARG* and *RARB*) from the homozygosity mapping and SNP concordance analyses suggests that, in addition to Wnt signaling, the retinoic acid signaling pathways also play a role in TH. The role of retinoic acid in limb development is controversial: while previous research has reported an association between hindlimb development and retinoic acid levels [23–25], a recent study found that hindlimb budding and patterning do not explicitly require retinoic acid signaling [26]. Our data support a role for retinoic acid receptor genes in hindlimb development as the severity of the TH phenotype appears to increase when mutations are present in these modifier genes. The data also support a role for Notch signaling, as *NOTCH1* was detected both in the SNP concordance analysis and the *de novo* assembly homozygosity mapping. *NOTCH1* and its ligand, *JAG2*, have repeatedly been implicated in hindlimb development [27,28].

We predict that modifier genes interact with *SMARCA4* and *WNT7A* and underlie the oligogenic inheritance pattern and subsequent variable phenotype. This hypothesis is supported by the structure of the network that was generated from the genes identified from the variant concordance mapping, GWAS, and homozygosity mapping analyses. However, the complex mode of inheritance of TH and the limited data available on both TH affected and control buffaloes preclude identification of the causal mutations, and the molecular mechanism by which the genes involved regulate hindlimb development remains unclear. The variants identified here in *SMARCA4* and *WNT7A* appear necessary for the expression of TH, but are located in introns based on computational predictions. It is possible that these variants are concordant with other variants on the same haplotype that are causal but were not detected during this study, due either to low sequence coverage or gaps in the reference assembly. For example, there are four gaps in *SMARCA4* in the current buffalo reference assembly and a large gap just upstream of the gene. The *WNT7A* gene is more complete, but also has one gap within the gene and three gaps in the upstream region. These gaps may contain genomic variants that alter the protein encoded by each gene or that alter gene expression through enhancers, promoters, or transcription factor binding sites. Likewise, the intronic variants identified may themselves disrupt unidentified regulatory elements. Variation in regulatory elements likely contributes to the expression of TH as severe reductions in the expression of *SMARCA4* in mice produce a similar phenotype [17].

We predict that selection against the variants found to be homozygous in *SMARCA4* and *WNT7A* in all cases but not in the controls and the avoidance of mating carriers of these variants would quickly eradicate the disease. This hypothesis could be tested by targeted genotyping of these loci in buffalo affected by TH and their parents to confirm that the loci are reliably predictive of the disease phenotype. The molecular dissection of the effects of mutations in *SMARCA4* and *WNT7A* could then be performed. However, this will likely require the collection of tissues from developing fetuses and possibly further development of the water buffalo reference genome and its annotation. Nevertheless, we hypothesize that eradication of the disease is now possible by selecting against the disease associated alleles for any of the concordant SNPs detected in *SMARCA4* and/or *WNT7A*.

## Methods

### Sample collection

DNA was collected from 4 bilaterally affected and 7 unilaterally affected TH cases and sequenced on an Illumina HiSeq 2500 at the Parco Tecnologico Padano in Milan, Italy. DNA sequences from 14 control buffaloes were provided by the International Water Buffalo Genome Consortium and were sequenced on an Illumina Genome Analyzer at the USDA Beltsville Research Center, USA. Although the pedigree of all sampled individuals was unknown, a principal component analysis conducted with smartPCA from the EIGENSOFT package [29] subsequent to variant calling indicated that the cases were not more related to one another than they were to the controls (Figure S3). Disease phenotypes were recorded for each case and a phenotype score was calculated based on the number of major distal bones present in each limb ranging from 0 (complete loss of both hindlimbs) to 10 (unaffected control) (Table S1).

### Genome sequencing

All animals were sequenced using Illumina technologies and 2 x 100 bp paired end libraries. Sequence depth varied from 5X to 15X average genome coverage, however, the cases were sequenced to an average depth of 12X and controls to only 7X (Table S1). Raw FASTQ sequences have been deposited at [INSERT LOCATION/ACCESSION NUMBERS].

### Genome alignment and variant detection

Raw sequences were trimmed for adaptors and quality using Trimmomatic-0.33 [30]. The reads were then aligned to the UMD_CASPUR_WB_2.0 water buffalo reference assembly (GCF_000471725.1) using the BWA-MEM algorithm, version 0.7.10-r789 [31]. Subsequently, we built a variant calling pipeline according to GATK Best Practices and optimized the pipeline
for a scaffold level reference genome [32–34]. The pipeline included duplicate removal using Picard (http://broadinstitute.github.io/picard), INDEL realignment, SNP and INDEL discovery using HaplotypeCaller, and genotype calling with GenotypeGVCFs. Base quality score recalibration and variant quality score recalibration were not performed due to the lack of availability of a known reference set of polymorphic sites in water buffalo.

SNP and INDEL variant sites were independently filtered. SNPs were filtered based on the number of detected alleles < 3 (biallelic), QD (Variant Confidence/Quality by Depth) < 2.0, FS (Phred-scaled *p*-value using Fisher’s exact test to detect strand bias) > 60.0, SOR (Symmetric Odds Ratio of 2×2 contingency table to detect strand bias) > 4.0, MQ (RMS Mapping Quality) < 40.0, MQRankSum (Z-score From Wilcoxon rank sum test of Alt vs. Ref read mapping qualities) < -12.5, or ReadPosRankSum (Z-score from Wilcoxon rank sum test of Alt vs. Ref read position bias) < -8.0. INDELs were filtered based on QD < 2.0, FS > 200.0, SOR > 10.0, ReadPosRankSum < -20.0, or InbreedingCoeff (Inbreeding coefficient as estimated from the genotype likelihoods per-sample when compared against the Hardy-Weinberg expectation) < - 0. 8. Furthermore, SNPs were filtered on an individual animal basis by setting genotypes with a Phred-scale genotype quality (GQ) < 10 to missing.

### Case versus control concordance analysis

Filtered SNPs and INDELs were analyzed for concordance on a case *versus* control basis. This involved sorting variants such that all cases were homozygous for an allele for which none of the controls were homozygous. A missing genotype among the cases caused the variant to be rejected from the analysis, but a missing genotype among the controls was ignored due to the lower mean sequence coverage for the controls.

### Homozygosity mapping by de novo assembly

Three *de novo* genome assemblies were carried out: unilaterally affected TH cases, bilaterally affected TH cases, and controls. Each assembly was initiated by pooling sequence reads from four individuals with the respective phenotype. The reads were assembled using MaSuRCA-3.1.3 using default parameters [35]. We used a negative binomial generalized linear model with the glm.nb function from the MASS package [36] to estimate the mean and dispersion parameter for the contig lengths produced from each assembly. A *p*-value was calculated for each contig to test the hypothesis that the contig was significantly greater in size than the mean, and the *p*-values corrected for multiple testing by estimating *q*-values [37]. As regions of the genome for which all of the pooled individuals are homozygous for a single haplotype can be assembled into large contigs (because the assembly graph does not fork), we extracted contigs that were significantly larger than average and those in the 99^th^ percentile for size from each assembly. These contigs were aligned to the UMD_CASPUR_WB_2.0 reference genome assembly and intersected with the water buffalo gene annotation. Finally, the lists of genes in the largest contigs produced from each assembly were compared. Genes within regions that were homozygous in all TH cases, but not in controls, were identified as candidates for risk of TH.

### Genome-wide association study (GWAS)

Given our initial uncertainty as to the mode of inheritance of TH, two GWAS analyses were run. The first was a mixed-model case *versus* control analysis while the second attempted to recover phenotypic information regarding disease severity by scoring the TH phenotypes according to the number of missing hindlimb bones, as previously described. Association tests for both models were performed using univariate linear mixed models and likelihood ratio tests implemented in GEMMA (version 0.94) with a centered genomic relationship matrix [38].

### Candidate region mapping and annotation

Variants identified by the concordance analysis and GWAS were intersected with the water buffalo gene annotation. *De novo* assembled contigs were aligned to the UMD_CASPUR_WB_2.0 water buffalo reference genome assembly using MUMmer3.23 [39]. The resulting reference positions were also compared with the water buffalo gene annotation.

Additionally, buffalo scaffolds having either a concordant SNP in the case *versus* control analysis or a significant GWAS association and the top 1% of *de novo* contigs were aligned to the *Bos taurus* UMD3.1 reference genome assembly using MUMmer3.23 [39]. This allowed us to interpret potential causal loci from the context of a genome as opposed to the 367,000+ unplaced scaffolds.

### Candidate gene ontology and network analysis

Candidate genes identified from the concordance analysis, GWAS, and homozygosity mapping were uploaded to the BovineMine warehouse [40] to search for the GO term(s) related to each gene. Similarly, BovineMine was also use to compare the list of candidate genes with the list of genes associated with the GO term “embryonic limb morphogenesis.” Network analyses were performed using GeneMANIA [41]. gProfileR version 0.6.1 [42,43] was used to conduct gene ontology enrichment analyses using genes identified from SNP concordance analyses, GWAS from binary and quantitative phenotypes, and homozygosity mapping by *de novo* assembly (99^th^ percentile analysis).

## Acknowledgements

JFT was supported by grants 2011-68004-30214, 2011-68004-30367, 2013-68004-20364, and 2015-67015-23183 from the USDA NIFA AFRI. JED was supported by grants M0-HAAS0027 and 2016-68004-24827 from the USDA NIFA.

